# Shared Neural Dynamics of Sentence-Level Language Processing in Verbally Proficient Autistic and Non-Autistic Adults

**DOI:** 10.64898/2026.11.20.768199

**Authors:** Benjamin T. Stanislav, Margot Mangnus, Daniel Stehr, Ashley G. Lewis, Peter Hagoort, Jana Bašnáková, Arjen Stolk

## Abstract

Autistic individuals often show atypical language development during childhood, yet many attain typical language proficiency by adulthood. This raises the question of whether the neural processes supporting language differ in autism once language has been acquired. Using electroencephalography (EEG), we compared verbally proficient autistic and non-autistic adults during rapid serial visual presentation of structured sentences and pseudoword lists. Sentence processing elicited oscillatory responses across theta, alpha, beta, and gamma frequency bands, with distinct spatiotemporal profiles and hemispheric lateralization most pronounced in the theta, alpha, and beta bands. Notably, beta-band activity increased over sentence position and was positively associated with verbal ability, consistent with a role in the maintenance and integration of linguistic information. These effects were highly comparable across groups, with similar patterns observed at both the individual and group levels and across sensor- and source-level analyses. Together, these findings indicate largely shared neural dynamics of sentence-level language processing in verbally proficient autistic and non-autistic adults, suggesting that atypical language-related neural organization is not an inevitable feature of autism in adulthood.

**Lay Summary:** Autistic individuals often experience difficulties during language development, but many become highly proficient language users as adults. Our findings show that verbally proficient autistic and non-autistic adults exhibit strikingly similar brain activity during sentence processing, indicating that atypical neural processing of language is not an inevitable feature of autism in adulthood.

## 1 Introduction

Although autism is often first recognized through atypical or delayed language development, language outcomes are highly heterogeneous. Approximately one third of individuals remain minimally verbal, while many others attain fluent and structurally typical language by adulthood (Helen Tager-Flusberg and Kasari 2013; Velikonja et al. 2019; Gagnon et al. 2021; Anderson et al. 2007; Short and Schopler 1988). This developmental heterogeneity raises the question of whether, among those who attain language proficiency, the neural processes supporting language nevertheless differ from those of non-autistic individuals.

Existing neuroimaging evidence has largely pointed toward altered language-related neural organization in autism, with both increased and reduced responses reported across regions implicated in linguistic function (Groen et al. 2008; Philip et al. 2012; Tryfon et al. 2018; Just et al. 2004; Jochaut et al. 2015). Among the most consistent findings is reduced left-hemisphere lateralization, observed across semantics, phonology, and sentence processing (Müller et al. 1999; Jouravlev et al. 2020; Dawson et al. 1986; Eyler et al. 2012; Coffey-Corina et al. 2008; Knaus et al. 2008; Lindell and Hudry 2013; Redcay and Courchesne 2008). Yet this pattern is not universal (Whitehouse and Bishop 2008), and much of the literature focuses on children, individuals with language delay, or samples with lower verbal ability, limiting conclusions about the mature language system in verbally proficient adults (Herringshaw et al. 2016). If these differences depend partly on developmental factors or ongoing language difficulties rather than on autism per se, the neural dynamics supporting language processing may be largely similar between verbally proficient autistic and non-autistic adults.

Testing this possibility requires characterizing not only where language-related activity occurs, but also how it unfolds as linguistic information is integrated over time (Nelson et al. 2017; Fedorenko et al. 2016; Piai et al. 2016). Electrophysiological approaches such as electroencephalography (EEG) are well suited to this goal, providing millisecond-level resolution of frequency-specific activity associated with the transient coordination of distributed neuronal populations into functional networks (Singer 1999; Varela et al. 2001; Bastos et al. 2014). During sentence processing, oscillatory activity emerges across theta (3-7 Hz), alpha (8-13 Hz), beta (14-20 Hz), and gamma (60-80 Hz) bands, with distinct spatial and temporal profiles linked to different aspects of linguistic computation (Bastiaansen and Hagoort 2015; Bastiaansen et al. 2010; Zioga et al. 2023; Terporten et al. 2019; Lam et al. 2016). In particular, beta-band activity increases across sentence position, a pattern linked to the maintenance and integration of linguistic information, though the precise computational role remains debated (Ashley Glen Lewis et al. 2023; Prystauka and Lewis 2019; Ashley G. Lewis and Bastiaansen 2015). Oscillatory lateralization is less well characterized, particularly in autism, but developmental work suggests that hemispheric asymmetry in alpha and beta, and to a lesser extent theta, changes over the course of language maturation (Nix et al. 2024; Spironelli and Angrilli 2010). Examining lateralization in these bands may therefore provide a complementary electrophysiological perspective on mature language organization (Olulade et al. 2020).

In the present study, we used EEG to characterize the neural dynamics of sentence-level language processing in verbally proficient autistic and closely matched non-autistic adults, examining how sentence-related oscillatory activity unfolds across time, frequency bands, and hemispheres. Participants viewed structured sentences and pseudoword sequences using rapid serial visual presentation (Fedorenko et al. 2010), which presents items sequentially at a fixed location, minimizing saccadic eye movements while allowing neural dynamics to be tracked as linguistic information accumulates over the course of a sentence (Rubin and Turano 1992). Pseudoword sequences preserved sequential presentation and position-dependent effects while disrupting the lexical-semantic and combinatorial information that supports coherent sentence processing (Fedorenko et al. 2020). A post-sequence probe assessed task engagement. Our primary analysis tested whether the resulting sentence-related oscillatory dynamics differed between autistic and non-autistic adults. Complementary analyses assessed their robustness at the individual level and characterized their hemispheric organization across sensor and source space.

## 2 Methods

### 2.1 Participants

A total of 42 autistic adults and 47 non-autistic participants were recruited through the Radboud University participant database, social media, and outpatient clinics. All participants were native Dutch speakers with normal hearing and normal or corrected-to-normal vision. Exclusion criteria included severe cognitive or neurological impairment, systemic disease, and current use of psychotropic medication or glucocorticoids. Autistic participants had received a clinical diagnosis of Autism Spectrum Disorder from a qualified clinician, which was confirmed via telephone screening. Four participants were excluded due to technical issues during EEG acquisition. Three additional participants were excluded prior to analysis to obtain groups closely matched on demographic and cognitive characteristics, yielding a final EEG sample of 41 autistic and 41 non-autistic participants (Table 1). Behavioral data from the memory probe task were available for 39 participants per group. The groups were matched on sex, age, handedness, and verbal and nonverbal IQ (Wechsler 1997; Raven 1989; Veale 2014), but differed significantly in Autism-Spectrum Quotient scores (Baron-Cohen et al. 2001). Both groups scored well above population averages on the Vocabulary and Similarities subscales of the Wechsler Adult Intelligence Scale. All participants provided written informed consent. The study was approved by the local ethics committee (Committee for Human-Subject Research, region Arnhem-Nijmegen; file no. 2019-6059), and participants received financial compensation.

**Table 1.**
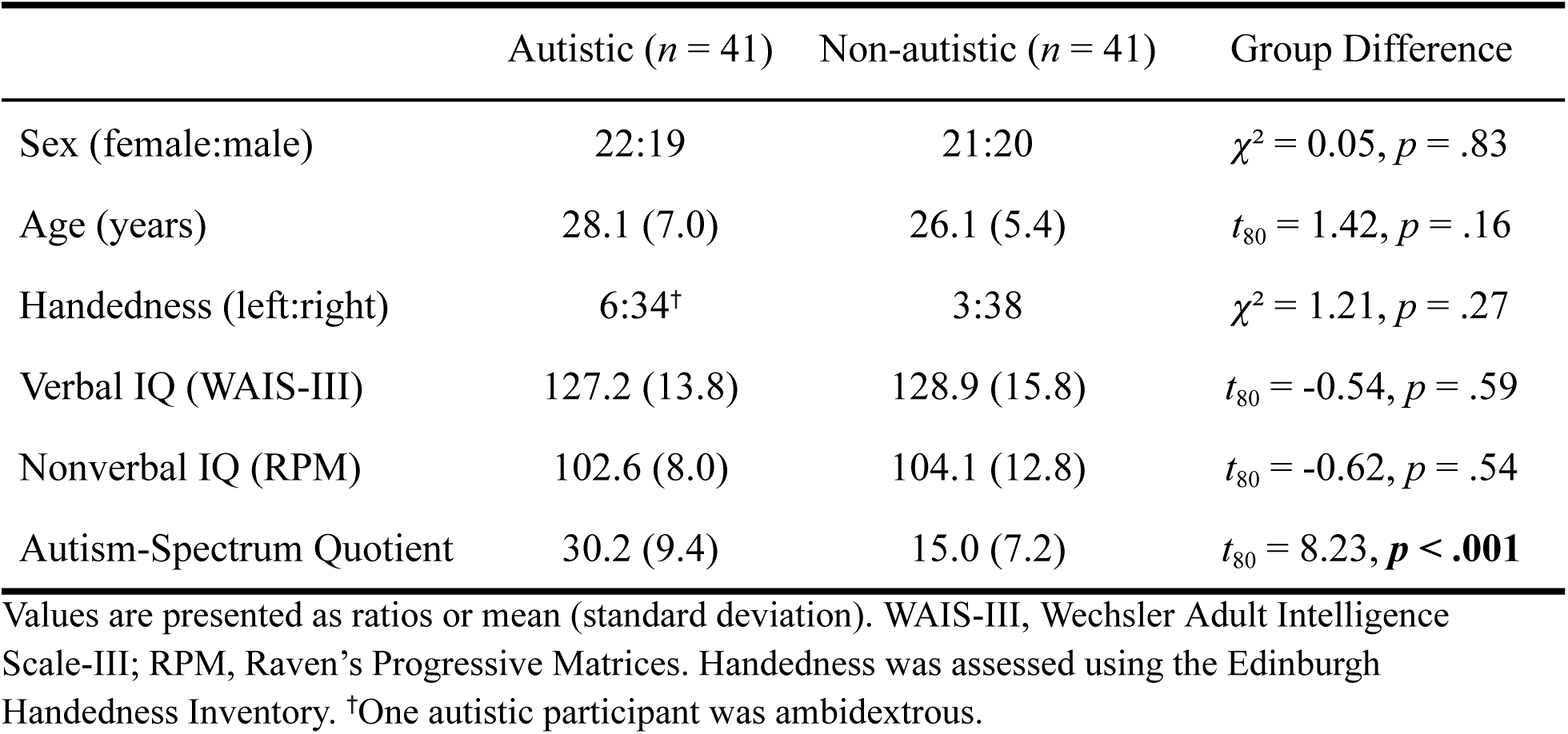
Demographic and cognitive characteristics.

|  | Autistic (*n* = 41) | Non-autistic (*n* = 41) | Group Difference |
|---|---|---|---|
| Sex (female:male) | 22:19 | 21:20 | *χ*² = 0.05, *p* = .83 |
| Age (years) | 28.1 (7.0) | 26.1 (5.4) | *t*<sub>80</sub> = 1.42, *p* = .16 |
| Handedness (left:right) | 6:34<sup>†</sup> | 3:38 | *χ*² = 1.21, *p* = .27 |
| Verbal IQ (WAIS-III) | 127.2 (13.8) | 128.9 (15.8) | *t*<sub>80</sub> = -0.54, *p* = .59 |
| Nonverbal IQ (RPM) | 102.6 (8.0) | 104.1 (12.8) | *t*<sub>80</sub> = -0.62, *p* = .54 |
| Autism-Spectrum Quotient | 30.2 (9.4) | 15.0 (7.2) | *t*<sub>80</sub> = 8.23, ***p* < .001** |

Values are presented as ratios or mean (standard deviation). WAIS-III, Wechsler Adult Intelligence Scale-III; RPM, Raven’s Progressive Matrices. Handedness was assessed using the Edinburgh Handedness Inventory. <sup>†</sup>One autistic participant was ambidextrous.

### 2.2 Experimental design

Participants read structured sentences and pseudoword sequences using a rapid serial visual presentation (RSVP) paradigm (Fedorenko et al. 2010). Each sequence consisted of 12 items presented sequentially at a fixed screen location (400 ms per item), supporting incremental processing while minimizing eye movements. Pseudoword sequences were constructed by replacing content words with pronounceable pseudowords derived from the original lexical items by recombining their constituent syllables, while preserving function words (Table 2). This manipulation preserved sequential presentation while disrupting sentence-level coherence, providing a contrast sensitive to the lexical-semantic and combinatorial processes that unfold during sentence processing (Fedorenko et al. 2020). Two independent native Dutch speakers verified that the pseudowords did not resemble existing Dutch words.

**Table 2.**
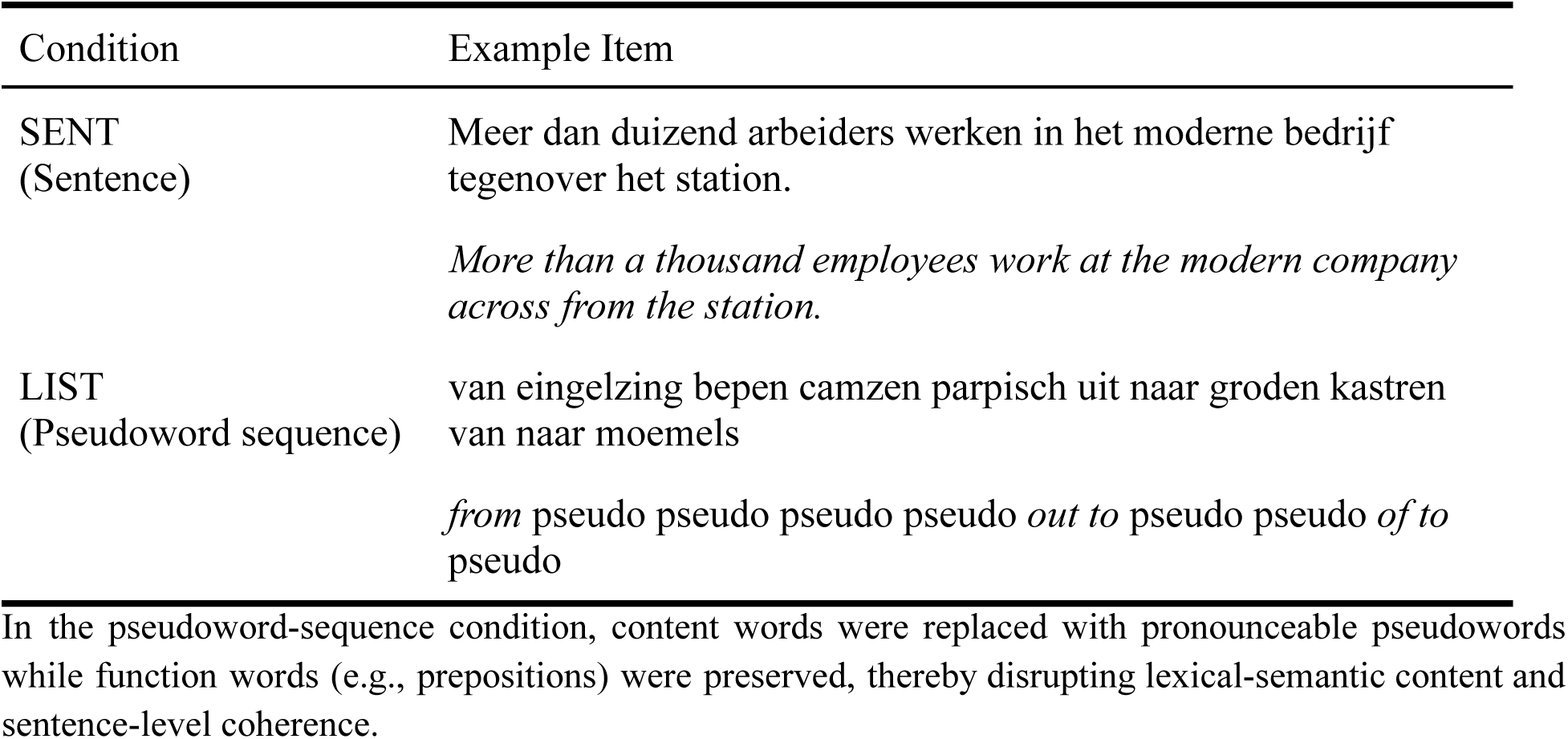
Example stimuli for the two experimental conditions with English translations.

| Condition | Example Item |
|---|---|
| SENT<br>(Sentence) | Meer dan duizend arbeiders werken in het moderne bedrijf tegenover het station. |
|  | *More than a thousand employees work at the modern company across from the station.* |
| LIST<br>(Pseudoword sequence) | van eingelzing bepen camzen parpisch uit naar groden kastren van naar moemels |
|  | *from* pseudo pseudo pseudo pseudo *out to* pseudo pseudo *of to* pseudo |

In the pseudoword-sequence condition, content words were replaced with pronounceable pseudowords while function words (e.g., prepositions) were preserved, thereby disrupting lexical-semantic content and sentence-level coherence.

Sentence topics varied across trials, and conditions were closely matched on several low-level properties, including total letter count per trial (sentences: M ± SD = 72.5 ± 5.8; lists: 72.6 ± 6.2), letter count per item (5.12 ± 2.64 vs. 5.12 ± 2.71), and syllable count per trial (20.7 ± 2.1 vs. 20.8 ± 2.2). Bigram frequency, derived from the SUBTLEX-NL database (Keuleers et al. 2010), was also comparable between conditions (6.04 ± 0.34 vs. 6.01 ± 0.37). Orthographic neighborhood density, quantified using OLD20—the mean orthographic Levenshtein distance to the 20 nearest real-word neighbors (Yarkoni et al. 2008)—differed between conditions (1.39 ± 0.63 vs. 1.71 ± 1.04; *t* = -6.46, *p* < .001), as expected from the lexical-form differences introduced by the pseudoword manipulation.

The experiment comprised 96 trials (48 per condition), presented across 32 three-trial blocks. Stimuli were randomized within condition and assigned to blocks, which alternated between pseudoword and sentence conditions, beginning with a pseudoword block. Each block began with a fixation cross of variable duration (4000-7750 ms), and each trial began with a 200-ms fixation cross followed by the 12-item sequence. To encourage attentive reading, participants then completed a memory probe indicating whether a specific word had appeared in the preceding sequence. The probe was displayed for 750 ms, although responses could be made after its offset; approximately two-thirds of responses occurred within the probe window, with no group differences in the proportion of responses made during versus after this window (Mann-Whitney U test, *p* = .16). Yes/No probes were balanced within each condition (25:23). Stimulus presentation and response collection were controlled using Presentation (Neurobehavioral Systems, Berkeley, CA, USA), which also delivered event triggers to the EEG recording system. Stimuli were presented centrally in black 18-point Courier New on a light gray background, with probe items and response options shown in red (Figure 1).

**Figure 1.**
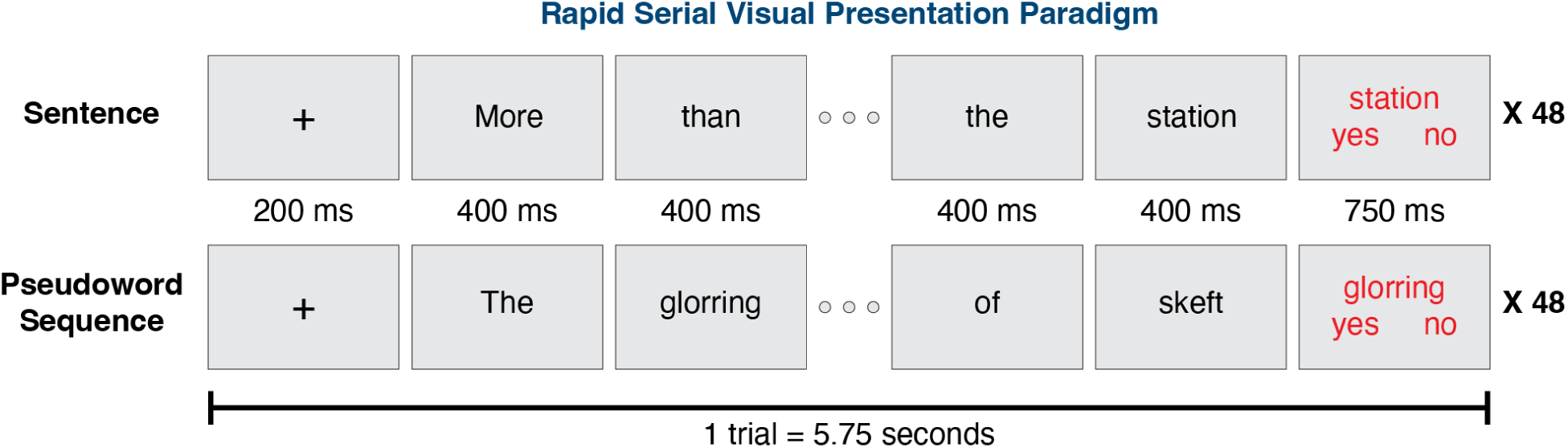
Rapid Serial Visual Presentation (RSVP) paradigm. Participants read structured sentences and pseudoword sequences presented word-by-word at a fixed screen location (400 ms per item) while EEG was recorded. In the pseudoword condition, content words were replaced with pronounceable pseudowords. Each trial began with a fixation cross, followed by a sequence of twelve words or pseudowords, and ended with a memory probe asking whether a specific word had appeared in the preceding sequence. Each condition comprised 48 trials.

### 2.3 Data acquisition

The experiment was conducted in an electrically shielded and sound-attenuated room. EEG was recorded from 60 electrodes using an actiCAP system with active sensors, BrainAmp DC amplifiers, and BrainVision Recorder software (Brain Products GmbH, Germany). Electrodes were positioned according to the extended international 10-20 system, with the left mastoid used as the online reference and AFz as the ground. Electrode impedances were maintained below 25 kΩ throughout the experiment. Signals were sampled at 1000 Hz and recorded with an online high-pass filter of 0.1 Hz; additional band-pass filtering and re-referencing were applied offline during preprocessing. Vertical eye movements were monitored using electrodes placed above and below the right eye, and horizontal movements were recorded using electrodes placed at the outer canthi of both eyes. To minimize head movement, participants’ heads were stabilized using a head-and-chin rest positioned 59 cm from the display screen.

To enable source-level analyses, electrode positions were obtained by performing a three-dimensional scan of each participant’s head while wearing the EEG cap using an iPad-based depth-sensing scanner (Structure Sensor; Occipital Inc., USA). The resulting head-surface model captured electrode locations and anatomical landmarks (nasion and left and right preauricular points), allowing co-registration with each participant’s anatomical MRI. High-resolution T1-weighted anatomical images were acquired for forward-model generation using a Siemens 3T MRI scanner with a magnetization-prepared rapid acquisition gradient-echo (MPRAGE) sequence (TR = 2200 ms, TI = 1100 ms, TE = 2.6 ms, flip angle = 11°, voxel size = 0.8 mm isotropic, acceleration factor = 2).

### 2.4 Behavioral analysis

Participant engagement and task performance were assessed using response accuracy and reaction times on the memory probe task. Reaction times were analyzed using generalized linear mixed-effects models in R (Bates et al. 2015). Fixed effects included condition (sentences, pseudoword lists), group (autistic, non-autistic), and their interaction, with by-participant random intercepts. Reaction times were modeled using an inverse Gaussian distribution with a log link (Lo and Andrews 2015). Accuracy was analyzed using a binomial mixed-effects model with a logit link function using the same fixed- and random-effects structure. Fixed effects were evaluated using Wald tests based on the model-estimated coefficients and standard errors. An additional mixed-effects logistic regression tested whether the condition effect on accuracy persisted after accounting for probe position on yes trials.

### 2.5 EEG preprocessing

Continuous EEG data were processed in MATLAB (R2024a) using FieldTrip (Oostenveld et al. 2011). Data were band-pass filtered between 1 and 150 Hz and notch-filtered at 50 and 100 Hz to attenuate line noise. Continuous recordings were visually inspected using FieldTrip’s summary and browser tools to identify noisy channels and artifactual segments. Noisy channels were removed and subsequently interpolated for sensor-level analyses only. EEG channels were re-referenced to the common average, and vertical and horizontal electrooculogram (EOG) signals were computed as bipolar derivations from periocular electrodes. Independent component analysis was applied to identify components reflecting ocular artifacts, which were removed based on their scalp topography and correspondence with the bipolar EOG signals. Data were then segmented into 400-ms epochs time-locked to each word presentation. After artifact rejection, an average of 0.13% (SD = 0.42%) of epochs was excluded. Exclusion rates did not differ significantly between groups or conditions (all *p*s > .39).

### 2.6 Time-frequency analysis

Each 400-ms word epoch was zero-padded to 1 s prior to Fourier transformation to increase frequency resolution (to 1 Hz) while preserving the effective temporal resolution of the 400-ms data window. Spectral power was computed using a Hanning taper for the theta (3-7 Hz), alpha (8-13 Hz), and beta (14-20 Hz) bands, and a Slepian multitaper with 5-Hz spectral smoothing for the gamma band (60-80 Hz). Frequency bands were selected based on prior work linking oscillatory dynamics to language processing (Bastiaansen and Hagoort 2015). Power values were *z*-scored within participants and averaged across frequency bins within each band. Analyses were restricted to words 2-12 of each sequence to minimize trial-onset effects.

At each channel and word position, mean oscillatory power across trials was analyzed using a mixed-design ANOVA with condition (sentences, pseudoword lists) as a within-subject factor and group (autistic, non-autistic) as a between-subjects factor. For cluster-based permutation testing, ANOVA-derived *F*-statistics were converted to signed *t*-equivalent values (√*F* with the sign of the corresponding effect) to preserve directional information during clustering. Samples exceeding a cluster-forming threshold of *p* < .05 were clustered across neighboring channels and adjacent word positions, with channel adjacency defined from the 60-channel electrode layout using FieldTrip’s triangulation method. Cluster-level statistics were calculated as the sum of the signed *t*-equivalent values within each cluster. Statistical significance was evaluated against a permutation distribution of the maximum cluster statistic generated from 5,000 permutations (Maris and Oostenveld 2007). Clusters falling within the upper or lower 2.5% of this distribution were considered significant (two-sided Monte Carlo *p* < .05). Permutation tests were performed separately for each frequency band and effect of interest.

For each significant cluster, the condition contrast was averaged across its constituent channels and word positions to yield a single cluster-level value per participant. These values were used both to quantify the magnitude of the condition effect using Cohen’s *d* with 95% confidence intervals derived from the noncentral *t* distribution (Meyer et al. 2021), and to assess evidence for the absence of group differences using Bayesian linear models. Bayes factors (BF_01_) compared models with and without group as a factor, with values greater than 1 favoring the null model and values between 3 and 10 typically interpreted as moderate evidence (Rouder and Morey 2012). To ensure stability, Bayes factor estimates were averaged across 100 iterations.

To characterize how oscillatory responses evolved over sentence position, peak electrodes were defined from the main effect of condition. For each frequency band, the electrode with the highest mean *t*-value across significant word positions and its four nearest neighbors defined a five-electrode cluster. Spectral power was averaged across these electrodes to obtain participant-specific time courses, and linear slopes across word positions were estimated for each participant. These slopes were entered into a multiple linear regression model to assess their relationship with individual differences in language ability. Verbal IQ served as the primary predictor, with group, autistic traits (AQ), and memory-probe accuracy included to account for diagnostic status, dimensional autistic traits, and individual differences in task performance, respectively. Variance inflation factors indicated low multicollinearity among predictors (all VIFs < 1.81). Based on prior work implicating beta-band activity in the maintenance and integration of linguistic information (Ashley Glen Lewis et al. 2023), this frequency band was of particular interest.

### 2.7 Decoding analysis

To assess whether sentence-level language processing could be detected at the individual level, linear support vector machine (SVM) classifiers were trained separately for each participant to distinguish sentence from pseudoword-list trials using *z*-scored spectral power averaged across word positions 2-12. Each trial was represented by a feature vector comprising 60 channels × 4 frequency bands. Classification performance was evaluated using repeated stratified 10-fold cross-validation (10 repeats), yielding a mean decoding accuracy for each participant. Mean accuracies were tested against chance (50%) within each group using one-sample *t*-tests and compared between groups using a two-sample *t*-test, with a Bayes factor (BF_01_) quantifying evidence for similarity between groups. Individual-level significance was assessed using permutation testing: condition labels were shuffled 1,000 times per participant, with the full cross-validation procedure repeated for each permutation to generate a participant-specific null distribution. Observed accuracies were compared against these distributions to obtain individual *p*-values, and the proportions of participants showing significant decoding (*p* < .05) were compared between groups using Fisher’s exact test.

### 2.8 Laterality index

To characterize the hemispheric organization of sentence-related oscillatory activity, participant-level condition contrasts were computed at each electrode as the difference between sentence and pseudoword-list trials, normalized by the pooled standard error across conditions. Laterality indices were then calculated for homologous left-right electrode pairs as the difference between left- and right-hemisphere condition effects (left - right), such that positive values indicate left-lateralized activity. These indices were analyzed using a mixed-design ANOVA with laterality (observed vs. zero) as a within-subject factor and group as a between-subjects factor. For significant lateralization clusters, effect sizes were calculated as described above to quantify both the magnitude of lateralization across participants and differences in lateralization between groups.

### 2.9 Source-level analysis

Source reconstruction was performed using Dynamic Imaging of Coherent Sources (DICS) beamforming (Gross et al. 2001), a frequency-domain approach that estimates source-level oscillatory power from cross-spectral density and forward models of volume conduction. A template grid in Montreal Neurological Institute (MNI) space (1 cm resolution) was warped to each participant’s anatomy to obtain subject-specific source grids. Volume conduction models were constructed using a three-shell boundary element method (Oostendorp and van Oosterom 1989), and electrode positions were co-registered to each participant’s MRI. Cross-spectral density matrices were estimated using a Slepian multitaper approach for each frequency band: theta (6 Hz ± 2 Hz smoothing), alpha (10 Hz ± 2 Hz), beta (17 Hz ± 3 Hz), and gamma (70 Hz ± 10 Hz). Frequency-specific spatial filters were computed using a regularization parameter set to 25% of the average sensor power (Ashley Glen Lewis et al. 2023) and subsequently applied to estimate source-level power at each grid point. Power estimates were *z*-scored and averaged across trials within each condition. Statistical analyses were conducted at each source location using the same mixed-design ANOVA and cluster-based permutation testing procedures as applied at the sensor level.

To assess source-level lateralization, participant-level condition contrasts were computed at each source location and mapped onto homologous locations across hemispheres. Laterality indices were then calculated as the difference between left- and right-hemisphere condition effects (left - right), such that positive values indicate left-lateralized activity, and were analyzed using the same mixed-design ANOVA framework as at the sensor level. In addition to whole-brain analyses, we performed a constrained analysis within a bilateral mask derived from significant lateralization clusters. Within this mask, source locations exceeding an uncorrected threshold (*p* < .01) for the condition contrast (two-sided *t*-test) were identified within each hemisphere for each participant. The number of suprathreshold locations was then quantified per hemisphere, providing a participant-level estimate of the spatial extent of sentence-related activity within the functionally defined region. Group-level statistics were subsequently computed on these participant-level counts. This approach complements cluster-based inference by capturing hemispheric differences in the spatial distribution of effects (Jouravlev et al. 2020).

## 3 Results

### 3.1 Behavioral performance

To verify task engagement and sensitivity to the stimulus manipulation, we examined performance on the memory probe task. Participants performed with high accuracy (81.5 ± 16.1%; M ± SD). Accuracy was higher for sentences (88.3 ± 15.8%) than for pseudoword lists (74.8 ± 13.3%; β = 1.52, SE = 0.12, *z* = 12.7, *p* < .001), consistent with the expected memory advantage for lexical and meaningful material (Baddeley et al. 2009; Massol et al. 2021). This condition effect remained after accounting for probe position (*OR* = 5.1, *p* < .001). Responses were also faster for sentences (661.2 ± 86.1 ms) than for pseudoword lists (758.5 ± 106.6 ms; β = -0.14, SE = 0.007, *z* = -19.1, *p* < .001). There were no significant group differences in accuracy (autistic: 82.4 ± 13.0%, non-autistic: 80.6 ± 18.7%; β = 0.11, SE = 0.21, *z* = 0.52, *p* = .60) or reaction time (autistic: 727.7 ± 111.9 ms, non-autistic: 692.0 ± 102.0 ms; β = 0.05, SE = 0.04, *z* = 1.30, *p* = .19), nor were there group-by-condition interactions for accuracy (β = -0.16, SE = 0.17, *z* = -0.96, *p* = .34) or reaction time (β = 0.002, SE = 0.01, *z* = 0.19, *p* = .85).

### 3.2 Oscillatory dynamics during sentence processing

We next examined how oscillatory dynamics differed between sentences and pseudoword lists. Consistent with prior work on language-related oscillations, sentences elicited greater spectral power than pseudoword lists across multiple frequency bands (Figure 2A). Significant condition effects were observed in the theta, alpha, and beta bands across most of the sequence (words 2-12; all *p*s < .001), with substantial effect sizes (theta: *d* = 0.88, 95% CI [0.63, 1.12]; alpha: *d* = 1.03 [0.77, 1.28]; beta: *d* = 1.16 [0.89, 1.43]). A condition effect was also observed in the gamma band later in the sequence (words 10-12; *p* = .033), although its magnitude was smaller (*d* = 0.26 [0.05, 0.47]).

**Figure 2.**
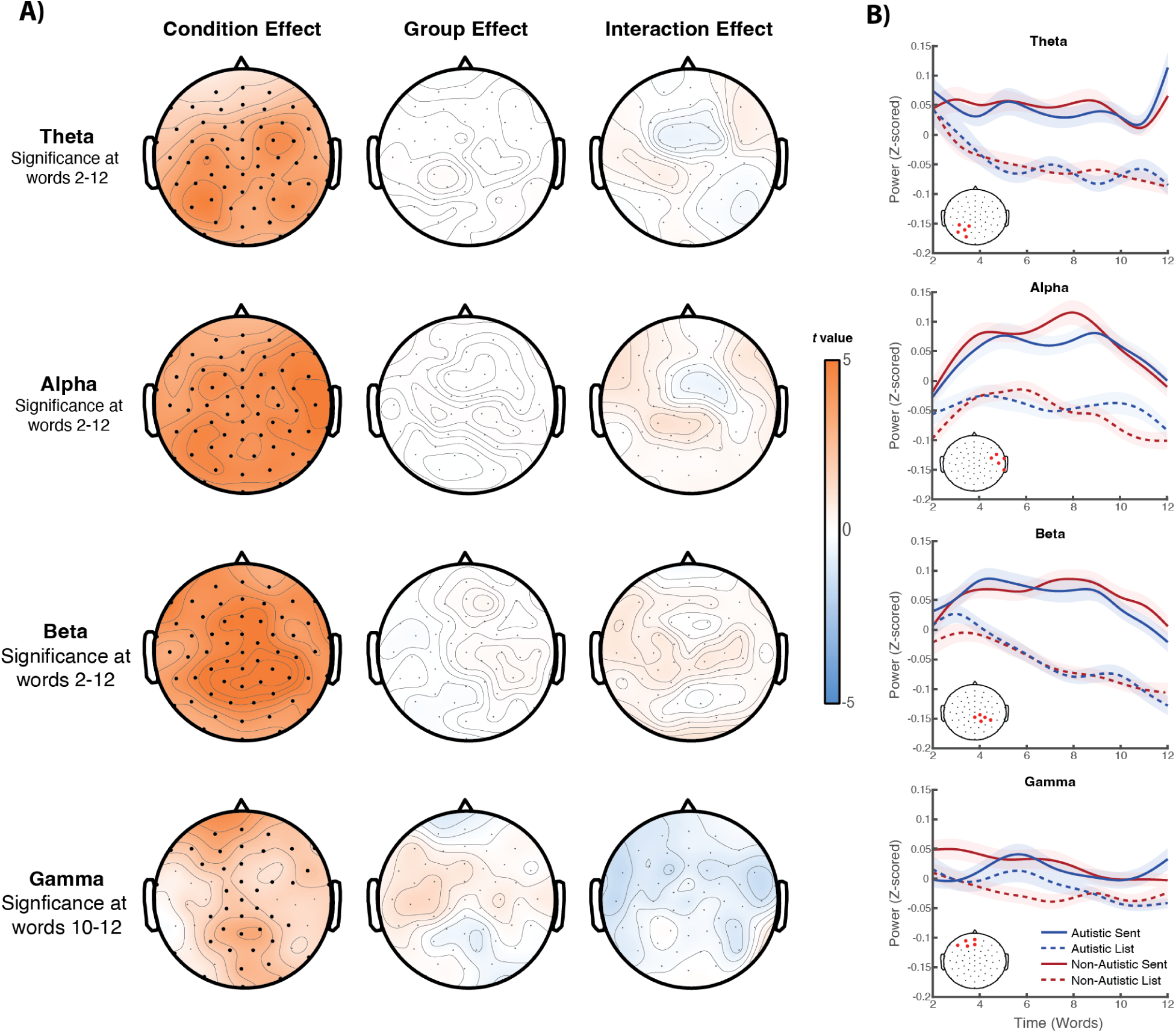
Oscillatory dynamics during sentence processing. (**A**) Topographic maps showing the main effects of condition (sentences vs. pseudoword lists), group (non-autistic, autistic), and their interaction across frequency bands. Colors represent *t*-values derived from cluster-based permutation tests, with electrodes belonging to significant clusters outlined. Significant condition effects were observed in the theta, alpha, and beta bands across words 2-12, and in the gamma band at later word positions. No significant group or interaction effects were observed. (**B**) Temporal evolution of spectral power averaged across electrodes exhibiting the strongest condition effects (indicated in the inset). Solid lines represent sentences and dashed lines pseudoword lists; blue denotes autistic and orange non-autistic participants. Shaded regions indicate ± SEM. Across both groups, spectral power diverged between conditions as sentences unfolded, consistent with incremental processing over time.

No clear group differences in these sentence-related effects were observed. Neither the main effect of group nor the group-by-condition interaction yielded significant clusters in any frequency band. Consistent with this, group differences within the condition-defined clusters were small, with confidence intervals spanning zero (theta: *d* = 0.07 [-0.36, 0.50]; alpha: *d* = 0.11 [-0.32, 0.55]; beta: *d* = 0.28 [-0.16, 0.71]; gamma: *d* = 0.09 [-0.34, 0.52]). Bayesian analyses provided converging evidence for the absence of group differences (theta: BF_01_ = 5.57; alpha: BF_01_ = 5.55; beta: BF_01_ = 5.53; gamma: BF_01_ = 5.23), further supporting similar sentence-related oscillatory responses in autistic and non-autistic participants.

To characterize how these responses evolved as linguistic information accumulated across the sentence, we next examined their temporal profiles, averaging spectral power across electrodes showing the strongest condition effects (Figure 2B). Across both groups, theta-, alpha-, and beta-band activity progressively diverged between conditions as sentences unfolded. The progressive increase in beta-band activity was of particular theoretical interest, given proposals linking beta oscillations to the maintenance and integration of linguistic information (Ashley Glen Lewis et al. 2023; Bastiaansen and Hagoort 2015). We therefore tested whether individual differences in beta slopes were associated with verbal ability independently of diagnostic status, autistic traits, and task performance. The overall regression model was significant (*F*(4,71) = 3.22, *p* = .017, adj. *R²* = .106), with verbal IQ emerging as the only significant predictor (*B* = 0.00029, *p* = .018). Higher verbal IQ was associated with a steeper increase in beta activity across sentence position, whereas group (*p* = .81), autistic traits (*p* = .054), and memory-probe accuracy (*p* = .92) were not significant predictors. No significant associations with verbal IQ were observed in the other frequency bands (all *p*s > .29).

### 3.3 Sentence decodability

The preceding analyses revealed robust sentence-related neural dynamics with no clear group differences. We next asked whether these effects were sufficiently robust to discriminate sentences from pseudoword lists at the level of individual participants. To this end, we trained a linear support vector machine classifier separately for each participant using trial-level spectral power across 60 channels and four frequency bands. Classification accuracy was significantly above chance in both groups (chance: 50%; autistic: 63.6 ± 8.7%, *t*(40) = 10.09, *p* < .001; non-autistic: 64.3 ± 9.6%, *t*(40) = 9.58, *p* < .001) and did not differ between groups (*t*(80) = 0.34, *p* = .73, BF_01_ = 4.13), providing further evidence that sentence-related neural signatures were similarly discriminable across groups. Within-participant permutation testing further showed significant decoding in the majority of participants in both groups (autistic: 28/41 (68%); non-autistic: 29/41 (71%)), with no difference in these proportions (*OR* = 1.12, *p* = 1.00).

### 3.4 Hemispheric lateralization

Beyond establishing sentence-related oscillatory effects, we next examined their hemispheric organization, given longstanding reports of altered language lateralization in autism. As shown in Figure 3, theta-band activity exhibited three spatiotemporal lateralization clusters, comprising right-lateralized anterior effects at words 7 and 12 (*p* = .010 and *p* = .038, respectively) and a left-lateralized posterior effect at word 12 (*p* = .008), although effect sizes were small across all three clusters (|*d*| ≤ 0.09). Alpha-band activity showed a sustained right-lateralized effect spanning words 8-12 (*p* < .001; *d* = -0.39, 95% CI [-0.60, -0.17]), while beta-band activity was right-lateralized at the final word (*p* = .006; *d* = -0.21, 95% CI [-0.42, 0.00]). No significant lateralization effects were observed in the gamma band.

**Figure 3.**
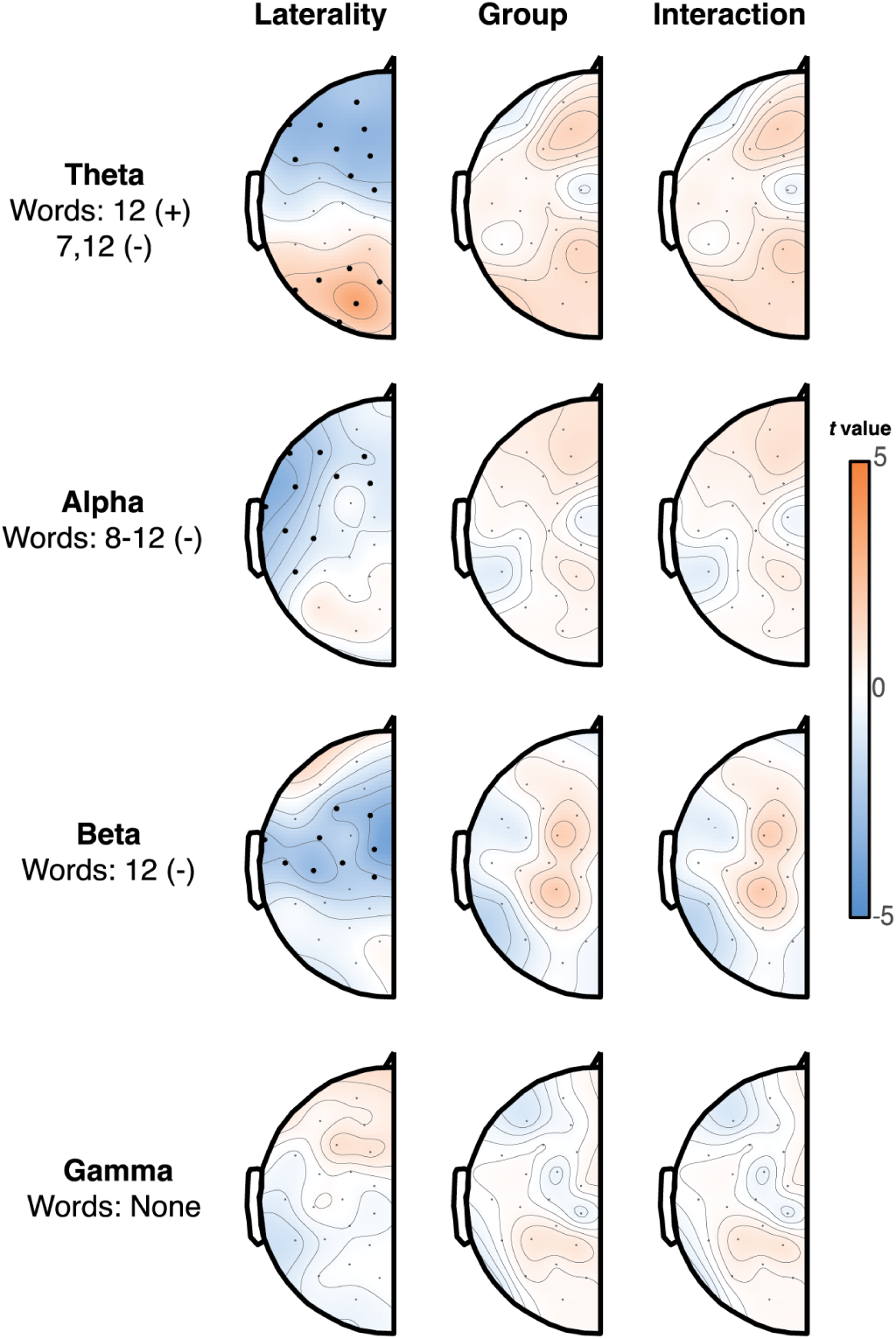
Hemispheric lateralization of sentence-related oscillatory activity. Topographic maps show lateralization indices (left - right) for the sentence-list contrast across frequency bands. Positive values indicate left-lateralization and negative values right-lateralization. Theta-band activity showed mixed lateralization at specific word positions, whereas alpha-band activity exhibited sustained right-lateralization (words 8-12) and beta-band activity showed right-lateralization at the final word. No significant lateralization was observed in the gamma band. No group or interaction effects were detected.

No clear group differences in these lateralization patterns were observed, with no significant main effects of group or group-by-laterality interactions in any frequency band. Within the lateralization-defined clusters, group differences were small and all confidence intervals included zero (alpha: *d* = 0.35, 95% CI [-0.09, 0.78], *p* = .12; beta: *d* = -0.17 [-0.60, 0.27], *p* = .45; theta: |*d*| = 0.07-0.40, with the largest difference observed for a right-lateralized cluster, *d* = 0.40, [-0.03, 0.84], *p* = .072). Bayesian analyses provided additional evidence favoring the absence of group differences for the alpha- and beta-band effects (BF_01_ = 3.39 and 4.35, respectively), with somewhat weaker to moderate evidence across the theta-band effects (BF_01_ = 2.91-4.28). Group effect sizes and Bayes factors were not calculated for gamma because no significant lateralization cluster was identified.

### 3.5 Source-level organization

To characterize the cortical organization underlying the sensor-level effects, we next examined their source-level distribution. Condition effects were widespread across the cortex in all frequency bands (all *p*s < .001), with no significant group differences or group-by-condition interactions. Hemispheric lateralization largely mirrored the sensor-level findings (Figure 4A). Theta-band activity showed an anterior right-lateralized effect (*p* = .007) and a posterior left-lateralized effect (*p* = .032), while alpha- and beta-band activity showed robust right-lateralization (both *p*s < .001), concentrated over inferior frontal and anterior temporal regions for alpha and more broadly over posterior and superior temporal regions for beta. No significant lateralization was observed in the gamma band.

**Figure 4.**
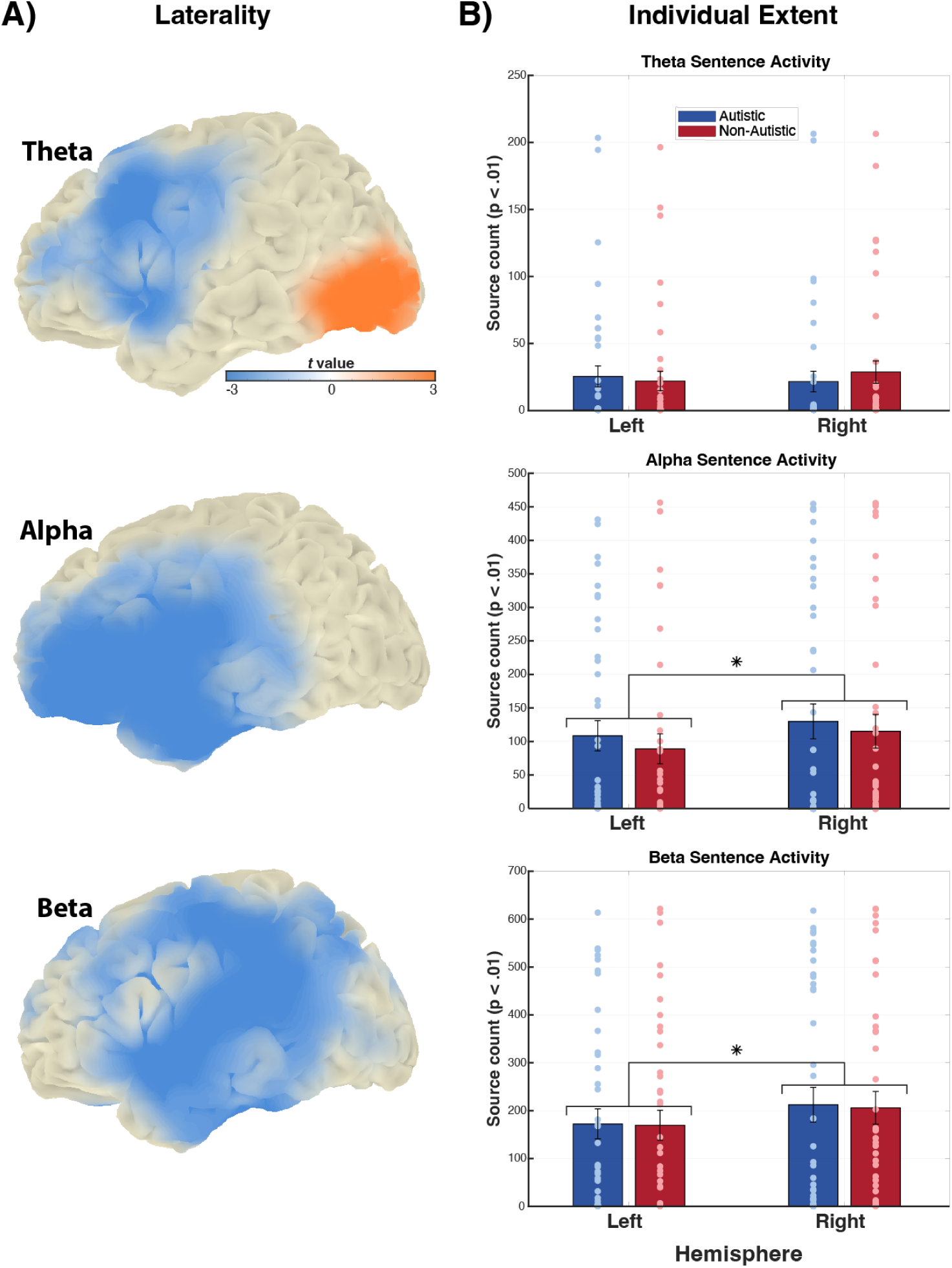
Source-level organization of sentence-related oscillatory activity. (**A**) Source-level maps of hemispheric lateralization for the sentence-list contrast across frequency bands. Positive values indicate left-lateralization and negative values right-lateralization. Consistent with sensor-level results, theta-band activity showed mixed anterior (right-lateralized) and posterior (left-lateralized) effects, whereas alpha- and beta-band activity exhibited predominantly right-lateralized patterns. (**B**) Participant-level quantification of hemispheric lateralization. Bars show the number of suprathreshold source locations (*p* < .01, uncorrected) within a bilateral mask derived from significant clusters, computed separately for each hemisphere and group. Points represent individual participants; bars indicate group means ± SEM; asterisks denote *p* < .05. Consistent with panel A, both groups showed the same overall pattern of hemispheric lateralization across frequency bands, with no significant group effects or interactions.

No significant group differences in these lateralization effects were observed (all *p*s > .13). Bayesian analyses provided moderate evidence favoring the absence of group differences for the theta and alpha effects (BF_01_ = 3.64-4.33), whereas evidence was weaker for beta (BF_01_ = 2.14).

To complement these group-level findings, we quantified the spatial extent of sentence-related activity within each hemisphere for individual participants, defined as the number of suprathreshold source locations within functionally defined group-level masks (Figure 4B). This analysis confirmed the posterior left-lateralized theta effect (*F*(1,80) = 5.64, *p* = .020) and right-lateralized alpha and beta effects (*F*(1,80) = 9.06, *p* = .004; *F*(1,80) = 8.91, *p* = .004, respectively), but not the anterior right-lateralized theta effect (*p* > .74). No group effects or group-by-laterality interactions were observed for these participant-level measures (all *p*s > .60), providing no evidence of systematic group differences in the hemispheric organization of sentence-related activity.

## 4 Discussion

Using EEG during sentence processing, we found largely shared neural dynamics in verbally proficient autistic and non-autistic adults. Across theta, alpha, beta, and gamma frequency bands, autistic and matched non-autistic participants showed highly similar oscillatory responses across sensor- and source-level analyses, with sentence-related activity also similarly discriminable at the individual level. Hemispheric organization likewise showed no clear group differences, while beta-band increases over sentence position, often linked to the maintenance and integration of linguistic information (Bastiaansen and Hagoort 2015; Ashley Glen Lewis et al. 2023), were associated with verbal ability but not diagnostic status. Together, these findings indicate that atypical language-related neural organization is not an inevitable feature of autism in adulthood.

At first glance, these findings appear to contrast with a neuroimaging literature that has largely pointed to altered language-related neural organization in autism, including differences in hemispheric lateralization (Groen et al. 2008; Tryfon et al. 2018; Herringshaw et al. 2016; Lindell and Hudry 2013). Much of this evidence, however, comes from studies of children or linguistically heterogeneous samples, in which developmental factors and ongoing language difficulties may contribute to observed differences. Studies of verbally proficient individuals have likewise reported similar neural responses between autistic and non-autistic participants during predictive parsing and semantic integration (Brennan et al. 2019; O’Rourke and Coderre 2021). Our findings extend this evidence to the temporal, spectral, and hemispheric organization of sentence-related neural activity. Differences in Autism have nevertheless been reported when linguistic integration places greater demands on resolving competing or conflicting information, as well as in relation to developmental language history (Ahtam et al. 2020; Peterson et al. 2024; Floris et al. 2021; Pijnacker et al. 2010; Enticott et al. 2013; Li et al. 2025). Together, this evidence suggests that neural differences in language processing may emerge under particular developmental and processing conditions rather than broadly characterizing autistic adults.

Importantly, similarity in sentence-level language processing does not imply an absence of broader differences in communication and context-sensitive processing (Boucher 2012). The present sample substantially overlapped with cohorts previously shown to exhibit such differences, including reduced responsiveness to interaction-based evidence about a communicative partner ((Koch et al. 2026); 71% overlap with the present sample) and more stereotyped neural responses during movie narratives ((Mangnus et al. 2024); 61% overlap). This distinction between language proficiency and communication is also apparent in development: autistic children matched to peers on language level nevertheless show differences in contingent discourse, particularly in building on a conversational partner’s preceding contribution (H. Tager-Flusberg and Anderson 1991). Thus, similar language abilities can coexist with differences in how language is integrated within an evolving communicative context.

These broader contextual processes may also contribute to heterogeneity in language acquisition. Early language learning depends not only on acquiring sensitivity to linguistic regularities, but also on embedding linguistic input within social and contextual frameworks that help render those regularities meaningful and learnable (Tomasello 2006; Kissine et al. 2023; Stolk et al. 2022; Stolk and Toni 2026; Jolliffe and Baron-Cohen 1999). Differences in these broader integrative processes could therefore shape language-development trajectories without implying corresponding differences in structural language processing among those who attain proficiency in adulthood. Once such structural regularities have been acquired, sentence-level processing may proceed similarly in minimally communicative contexts such as the present paradigm.

Beyond the question of group differences, the observed lateralization effects provide a complementary perspective on the hemispheric organization of neural dynamics during mature language processing. Alpha lateralization was most pronounced over inferior frontal and anterior temporal regions, whereas beta lateralization was more prominent over posterior and peri-sylvian regions, consistent with distinct roles of these frequency bands within language networks (Schoffelen et al. 2017). Notably, these effects arose primarily from sentence-related changes in right-hemisphere activity relative to the control condition (Figure 4B), consistent with developmental evidence for increased rightward lateralization of alpha and beta activity from childhood to adulthood (Nix et al. 2024; Spironelli and Angrilli 2010). This frequency-specific organization complements the predominantly left-lateralized language organization typically observed with fMRI, highlighting an additional dimension of hemispheric specialization captured by oscillatory activity.

The present findings should be interpreted in light of several limitations. First, the cross-sectional design cannot establish how the adult neural patterns observed here emerged over development, and participants’ early language-development histories were not available. Longitudinal studies will be needed to determine how language-related neural organization changes across development and relates to adult outcomes. Second, while the RSVP paradigm provides a broad contrast encompassing multiple linguistic processes, it examines sentence processing within a controlled, incremental setting rather than language use in richer communicative contexts, where group differences may be more pronounced or qualitatively different (H. Tager-Flusberg and Anderson 1991; Wadge et al. 2019). Extending this work across development and into more interactive contexts will be important for establishing the conditions under which the similarities observed here persist or give way to differences.

In conclusion, we found that sentence-level language processing engages largely shared neural dynamics in verbally proficient autistic and non-autistic adults. Across frequency bands, sentence-related oscillatory responses showed similar temporal and hemispheric organization, while beta-band increases, often linked to linguistic integration, were better explained by verbal ability than by autism diagnosis. These findings suggest that atypical language-related neural organization is not an inevitable feature of autism in adulthood. Together with prior evidence of communicative differences in this population, they underscore the need to distinguish how proficient adults process language from how language is acquired and used in communicative contexts.

## Author Contributions

BTS and MM conducted the analysis, interpreted the findings, and drafted the original manuscript. JB contributed to study design. AS contributed to data interpretation and drafting of the original manuscript. All authors contributed to the interpretation of the findings and preparation of the final manuscript.

## Acknowledgments

We would like to thank the study participants for their time and contribution to this research, and Sophia Walker and Daniela Santana Gutiérrez for their assistance with data acquisition.

## Ethics Statement

The study was approved by the Committee on Research Involving Human Subjects (Arnhem-Nijmegen region, the Netherlands).

## Informed Consent

All participants provided informed consent before taking part in the study

## Data Availability Statement

Data and analysis scripts supporting the findings of this study will be made publicly available via the Open Science Framework upon publication.

